# Multi-Timescale Neural Adaptation Failure as a Mechanistic Signature of Major Depression

**DOI:** 10.64898/2025.12.16.694510

**Authors:** Yunkai Yang, Yanli Zhao, Kun Xie, Chenfan Song, Yilun Huang, Anhui Kong, Shiqi Huang, Zixuan Xing, Wenyi Wang, Nils Muhlert, Rebecca Elliott, Shuping Tan, Zhenhong He, Dandan Zhang

## Abstract

Major depression (MD) is usually characterized in terms of momentary abnormalities, such as negative bias or emotional blunting, derived from paradigms that measure only the first response to a stimulus. Yet depressive experience in daily life is repetitive and cumulative. Here we test the hypothesis that a failure of neural adaptation across repetitions is a core feature of MD. Using a repeated negative-stimulus paradigm, we quantified adaptation at micro (within-trial) and macro (across-session) scales via electroencephalography in 28 MD patients and 34 healthy controls, measuring both conventional event-related potentials and large-scale traveling-wave propagation. MD patients exhibited pervasive maladaptation: neural rigidity at the micro scale (absent repetition suppression) and progressive regulatory depletion at the macro scale (escalating emotional reactivity with declining cognitive control). Across analyses, adaptation-based metrics revealed larger group differences than single-shot indices, and classifiers built on adaptation features—especially traveling-wave dynamics—outperformed single-shot models (task AUC = 0.83 vs 0.78) and generalized to two independent datasets (up to AUC = 0.91). These findings challenge conventional symptom-based characterizations of MD, establishing adaptation failure—not initial reactivity—as the mechanistically primary pathological signature. This reframing provides a unified mechanistic explanation for rumination and cognitive inflexibility while offering a standardized, low-cost biomarker requiring minimal electrodes (≤8 channels), enabling large-scale screening in resource-limited settings.

## 1 Introduction

Imagine this scenario: a beloved pet is lost, a wallet goes missing, or a close relationship breaks down. The first time it happens, it hurts. But when similar negative events happen again and again, most people gradually become “numb,” with the emotional sting fading over repetitions. This attenuation is not indifference but a protective form of neural adaptation that helps prevent excessive depletion of emotional resources under sustained adversity (Barthel et al., 2025; McEwen, 2000). It reflects experience-dependent neural plasticity: repeated exposure typically engages fast inhibitory reweighting and slower homeostatic processes that stabilize network excitability over time (Turrigiano, 2012). Patients with major depression (MD), however, often appear to lack this safeguard: each repetition feels as painful as the first, sometimes even more overwhelming (Gou et al., 2025; Klug et al., 2024).

Converging clues point to adaptation failure as a core feature of MD, yet systematic experimental verification remains lacking. Behaviorally, patients show persistent negative responses despite repeated exposures, reflected in heightened rumination (Marroquín & Nolen-Hoeksema, 2015), impaired habituation to emotional stressors (Siegle et al., 2002), and maladaptive regulatory patterns (Stange et al., 2017). Neuroscientific evidence converges on the same conclusion: persistent prefrontal–hippocampal and prefrontal-ventral striatum dysconnectivity during habituation (Disner et al., 2011; Heller, 2016; Liu et al., 2017; Tura & Goya-Maldonado, 2023), failure to downregulate amygdala activity across repetitions (Klug et al., 2024), aberrant striatal prediction-error signaling (Gradin et al., 2011; Kumar et al., 2008), and sustained gamma activity to repeated negativity (Siegle et al., 2010). This “loss of adaptive protection” has even been proposed as one of the core causal mechanism (Durisko et al., 2015) and is consistent with the DSM-5 emphasis on persistent negative affect in MD (American Psychiatric Association, 2022). Yet despite this converging but fragmented evidence, adaptation failure has never been directly quantified within a paradigm that reveals its temporal dynamics.

Existing accounts of MD responses to negative events—most prominently negative bias (heightened initial reactivity; (Elliott et al., 2002; Gotlib & Joormann, 2010) and emotional blunting (reduced overall responsiveness; Christensen et al., 2022)—have largely been treated as distinct phenomena from adaptation failure described above. We propose, however, that these accounts may be reconciled through a unifying possibility: negative bias or emotional blunting may represent specific temporal snapshots extracted from an underlying maladaptive trajectory—one that begins abnormally and remains so (or even worsens), rather than one that normalizes appropriately over time. If true, the apparent divergence between these accounts reflects not separate pathological features but different measurement windows onto the same dysfunctional process.

Critically, testing this trajectory-based account requires methods that are sensitive to temporal dynamics. Yet nearly all experimental work on MD relies on single-shot designs, in which negative stimuli are presented once (or sparsely intermixed with other stimuli) and responses are averaged as if each event were independent. While methodologically convenient, such designs can only reveal whether MD patients react more strongly or more weakly at the *initial* encounter; they cannot show whether these responses persist, fade, or escalate across repetitions. As a result, single-shot paradigms cannot distinguish elevated-but-adaptive reactivity from genuine adaptation failure, because they capture only the first snapshot of what may be a much longer maladaptive trajectory. This limitation is not merely technical but conceptual. Real-world depressive experiences are inherently repetitive and cumulative, encompassing recurrent stressors, sustained interpersonal criticism, and iterative emotional challenges (Kessler, 1997). Likewise, the DSM-5 defines MD not by momentary reactions but by prolonged negative states that fail to resolve over time (American Psychiatric Association, 2022). By sampling only isolated moments, single-shot designs fundamentally mismatch the temporal structure of MD psychopathology and risk obscuring the very processes—progressive resource depletion, impaired compensatory regulation, and breakdown of adaptive mechanisms—that emerge under sustained emotional stress (Mather & Sutherland, 2011). Directly testing the adaptation-failure hypothesis therefore requires quantifying the full temporal trajectory rather than isolated snapshots.

To address this gap, we employed a repeated negative-stimulus paradigm designed to approximate the progressive accumulation of emotional burden in MD. This design allowed us to recover conventional indices derived from single-shot paradigms while simultaneously quantifying adaptation at two complementary temporal scales. At the micro scale, we measured *within-trial repetition suppression*—the change from the first to the second viewing of the same stimulus (Garrido et al., 2009). At the macro scale, we characterized *habituation trajectories* across the full 20–30 min session, capturing how neural responses evolve cumulatively over repeated exposures (schematized in **Figure 1A**). To span levels of neural organization, we extracted both standard electroencephalographic (EEG) measures (N2, late positive potential (LPP), theta rhythms; Folstein & Van Petten, 2008; Kaiser et al., 2003) and large-scale travelling-wave propagation metrics (Muller et al., 2018), enabling us to link local electrophysiological responses to system-level spatiotemporal coordination.

**Figure 1.**
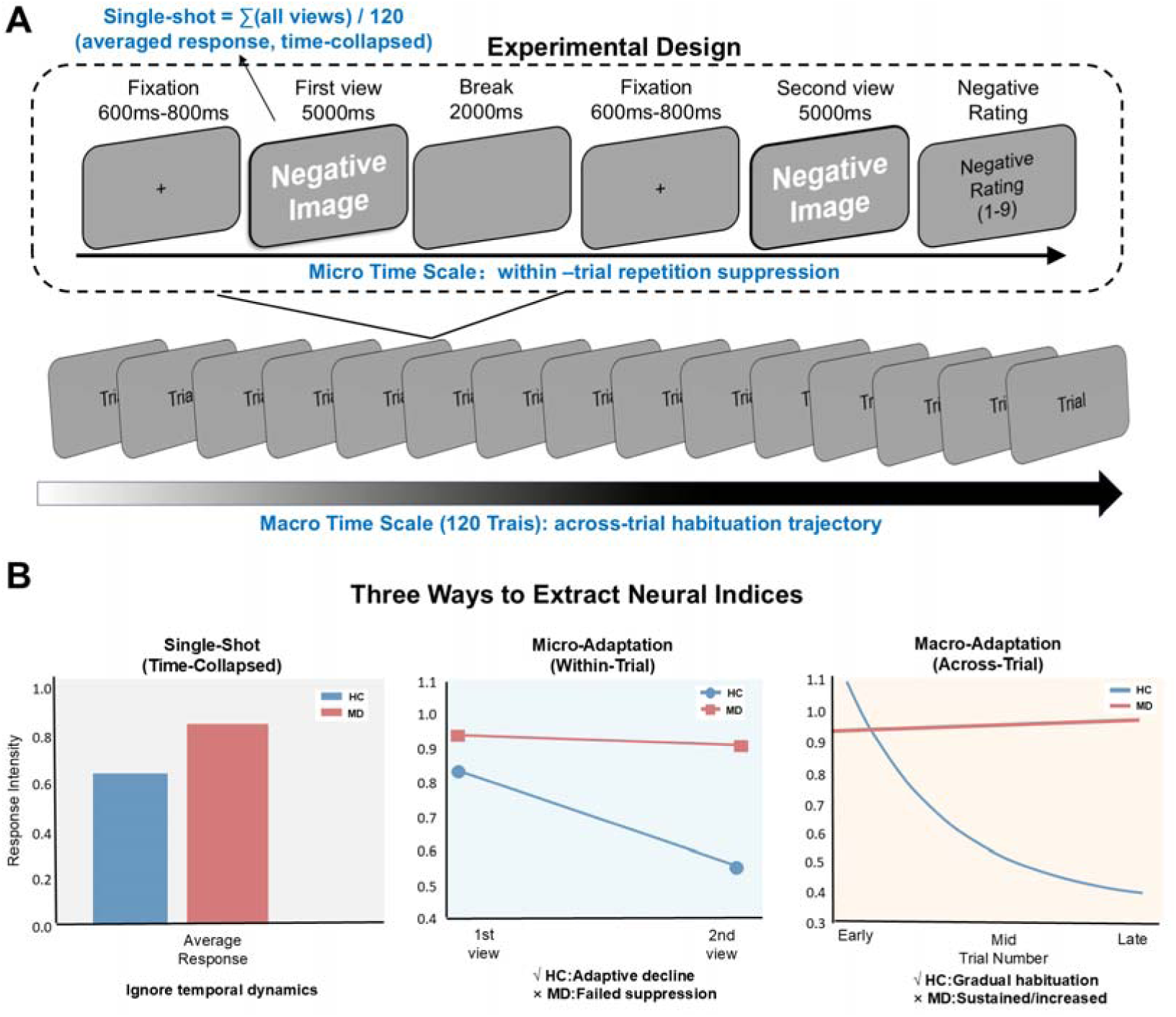
Experimental Design and Extraction of Multi-Scale Neural Indices. **(A) Paired-Presentation Paradigm.** Each trial consisted of a First View (5000 ms) and a Second View (5000 ms) of the same negative stimulus, separated by a 2000 ms break. This design enables the quantification of neural dynamics at two distinct timescales: the Micro Time Scale, indexing immediate repetition suppression within a trial, and the Macro Time Scale, indexing cumulative habituation across 120 trials. Images shown are schematic illustrations for conceptual purposes only and do not depict actual experimental stimuli or identifiable individuals. The traditional “Single-shot” metric (blue text) represents the averaged response across views, effectively collapsing the temporal dimension. **(B) Three Approaches to Extract Neural Indices.** Left (Single-Shot): A static measure of average response intensity that ignores temporal dynamics, often masking adaptation deficits. Middle (Micro-Adaptation): Quantifies the immediate change from the first to the second view. Healthy controls (HC) typically exhibit an adaptive decline (repetition suppression), whereas patients with Major Depression (MD) show failed suppression. Right (Macro-Adaptation): Tracks the cumulative trajectory from early to late trials. HCs show gradual habituation, while MD patients exhibit sustained or increased reactivity (sensitization/depletion) over the session.

Importantly, each temporal scale yielded two distinct classes of indices: (1) Single-shot indices, reflecting *initial reactivity* (e.g., the first-view N2, LPP, theta activity); (2) Adaptation indices, reflecting *dynamic change*—micro-scale slopes (first→second-view change in N2, LPP, theta, and travelling-wave velocity) and macro-scale slopes (N2, LPP, theta, and wave-velocity changes across the whole session; see **Figure 1B** for a schematic overview).If adaptation failure is mechanistically more central to MD than heightened initial reactivity, then adaptation indices should outperform single-shot indices both in forward comparisons (larger group differences) and reverse comparisons (superior diagnostic classification).

Our findings reveal a dual maladaptation profile in MD: neural rigidity at the micro-scale (failed repetition suppression) and progressive regulatory depletion at the macro-scale (absent or reversed habituation trajectories). Critically, adaptation-based measures revealed larger group differences and stronger classification performance than traditional single-shot indices. Whereas single-shot responses provided only moderate discrimination (area under the ROC curve (AUC) = 0.78), macro-scale adaptation dynamics—quantifying cumulative habituation trajectories—performed better (AUC = 0.80), and combining micro- and macro-level adaptation yielded the strongest task-state performance (AUC = 0.83). Feature-importance analyses further showed that traveling-wave parameters contributed the majority of discriminative information (59%), with backward waves (BW) contributing 3.1× more diagnostic variance than forward waves (FW) and wave velocity outweighing power by 2.2×, consistent with impaired top-down regulatory coordination in MD. Critically, this advantage was not task-specific: the same propagation signatures generalized robustly to two independent datasets—including a self-collected clinical cohort (AUC = 0.91) and a public resting-state dataset from Mumtaz et al., (2017; AUC = 0.82)—demonstrating that the adaptation signal reflects a stable, trait-like neural fingerprint of MD. Together, the dual-scale breakdown of adaptation gives a mechanistically account for why depressive affect remains persistently high despite repeated exposure, and produces a trait-like neural signature that outperforms single-shot indices in distinguishing MD from healthy functioning.

## 2 Results

### 2.1 Forward statistical inference

#### 2.1.1 Single-shot baseline neural responses to negative stimuli

To begin with, we examined how participants responded in the single-shot condition where negative stimuli were viewed for the first time (**SI Text 1**). We observed an unexpected pattern in the group comparison. The overall more negative emotional ratings observed in MD patients (*p* = 0.028, *Cohen’s d* = 1.18) support a generalized negative bias in this population (Gotlib & Joormann, 2010). At the event-related potential (ERP) level (**SI Text 2 and Figure S1**), MD patients demonstrated a marked reduction in the amplitudes of the later frontal LPP component compared with HCs (*p* = 0.029, *Cohen’s d* = −0.59). This indicates a significant emotional blunting occurred (Christensen et al., 2022).

These ERP findings were paralleled by the results at the oscillatory level (**SI Text 3**). We identified a significant elevation of the theta activity in overall PSD (*p <* 0.001, *Cohen’s d* = 1.12). Besides, traveling wave analyses further revealed an abnormal pattern of large-scale cortical propagation energy (**SI Text 5**): MD patients showed weakened FW theta energy (*p* = 0.012, *Cohen’s d* = −0.68), enhanced low beta-band energy (*p* = 0.019, *Cohen’s d* = 0.63) and high beta-band energy (*p* < 0.001, *Cohen’s d* = 0.10). Similarly, for BW of MD patients (**SI Text 6**), there is a higher alpha velocity (*p* = 0.012 *Cohen’s d* = 0.64). These characteristic signatures indicated a systematic shift in cross-regional coordination and signal propagation path of MD patient brains.

Taken together, the single-shot results indicated that MD patients were already in an abnormal baseline neural state, characterized by attenuated frontal LPP responses, globally increased low-frequency oscillatory power, and atypical cross-regional propagation dynamics. This pattern was highly consistent with prior evidence linking MD patients to emotional blunting, attentional engagement with negative information, and disruptions in large-scale neural network connectivity (Christensen et al., 2022; Gotlib & Joormann, 2010; Greicius et al., 2007).

#### 2.1.2 Micro-scale failure of fast neural adaptation

At a micro timescale, which was within the few seconds between the first and second view of the same negative image in a trial, we identified a robust failure of neural adaptation to negative stimulus repetition in MD patients (**SI Text 2**). Specifically, when the same negative image was presented for the second time, HCs showed the typical repetition suppression effect of the late ERP component, with frontal LPP amplitudes exhibiting marked suppression (*p* < 0.001, *Cohen’s d* = −0.64; **Figure 2A-B**). In contrast, the frontal LPP of MD patients had nearly no change from the first to the second viewing, producing a reliable group-by-time interaction (*p* =0.004, *η^2^p* = 0.13). This pattern indicated that upon the reappearance of an emotional stimulus, HCs rapidly down-regulated the neural load associated with emotional processing, whereas depressed brains fail to achieve such second-scale deactivation.

**Figure 2.**
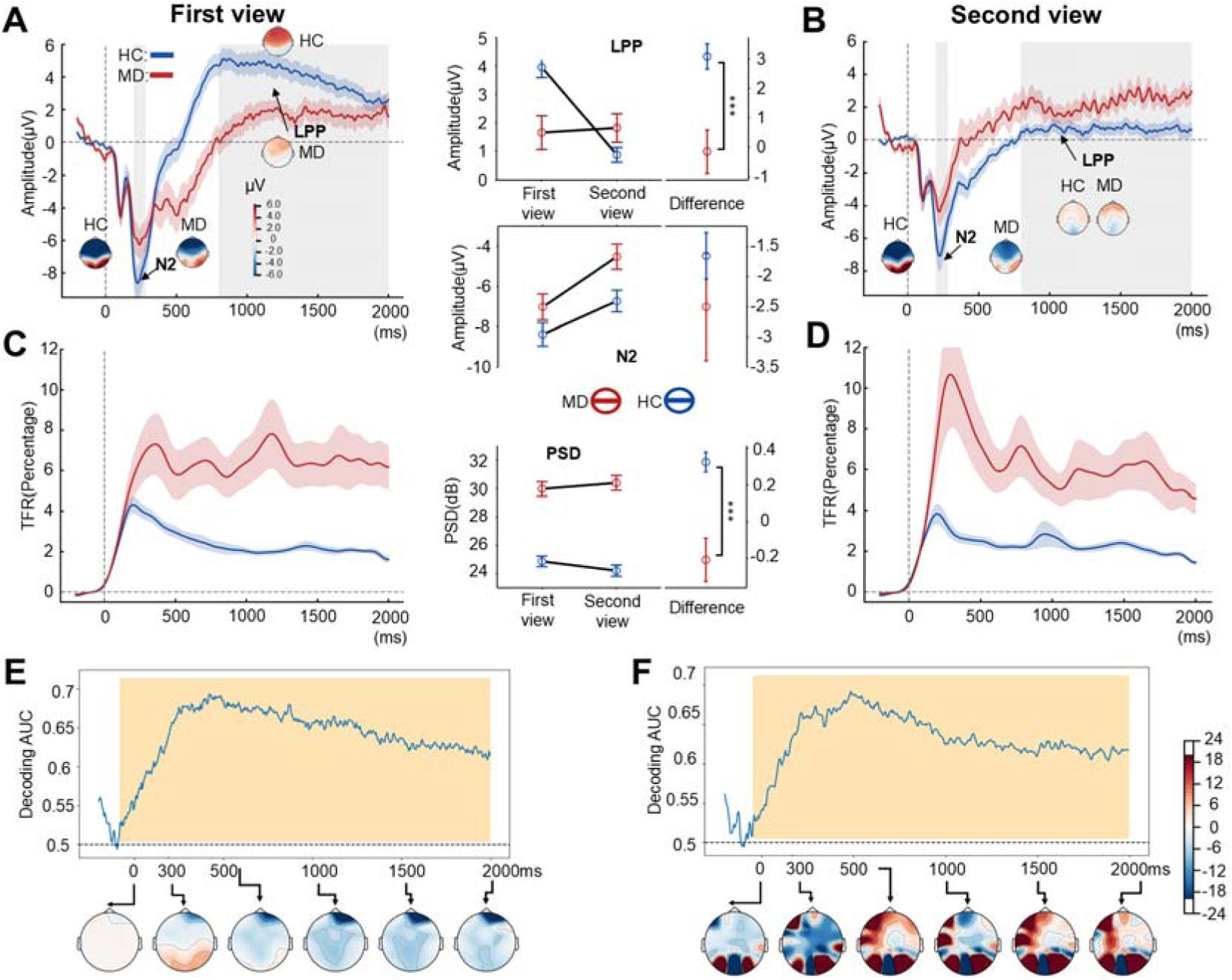
Micro-scale adaptation failure: Neural rigidity to immediate repetition. **(A-B) Grand average Event-Related Potential (ERP) at frontal electrodes during the first and second viewing phases.** Healthy controls (HCs; blue) display a robust repetition suppression effect, evidenced by a significant reduction in late positive potential (LPP) amplitude during the second view. Major depression (MD) patients (red) exhibit neural rigidity, with LPP amplitudes failing to attenuate. The central line charts quantify the mean amplitudes of N2, LPP components, and theta Power Spectral Density (PSD). **(C** – **D) Time-Frequency Representations (TFR) of frontal theta power.** HCs show rapid downregulation of theta activity upon stimulus repetition, reflecting efficient resource conservation. Conversely, MD patients maintain elevated theta engagement during the second view, indicating a failure to disengage cognitive control resources. **(E** – **F) Multivariate pattern analysis (MVPA) temporal generalization.** Decoding accuracy time-courses reliably distinguish first from second views in both groups. However, Haufe-transformed activation patterns reveal divergent mechanisms: Panel E shows that HCs exhibit global downregulation (negative weights), whereas Panel F shows that MD patients exhibit sustained or potentiated activation patterns (positive weights), confirming the absence of rapid neural adaptation. Shaded areas indicate standard error. *** *p* < 0.001.

In addition, pattern classification using time-domain multivariate pattern analysis (MVPA) further strengthened the evidence for micro-scale neural rigidity (**SI Text 4; Figure 2E-F**): although both groups could reliably distinguish the first from the second view (*p* < 0.05, cluster-corrected), the generally negative decoding weights in HCs reflected a weakening of the overall activation pattern, whereas the decoding weights in MD patients tended to be positive, indicating a tendency to maintain and even increase the activation evoked by the first view.

A consistent response was observed at the oscillatory level (**SI Text 3; Figure 3C-D**). We found that HCs showed a marked decrease in PSD of the theta power (*p* = 0.011, *Cohen’s d* = −0.39) during the second view time, revealing a typical rapid neural inhibition. Conversely, the theta activity in MD patients remained at a high level following repetition, constituting a characteristic signature of sustained resource engagement (Cavanagh & Frank, 2014).

**Figure 3.**
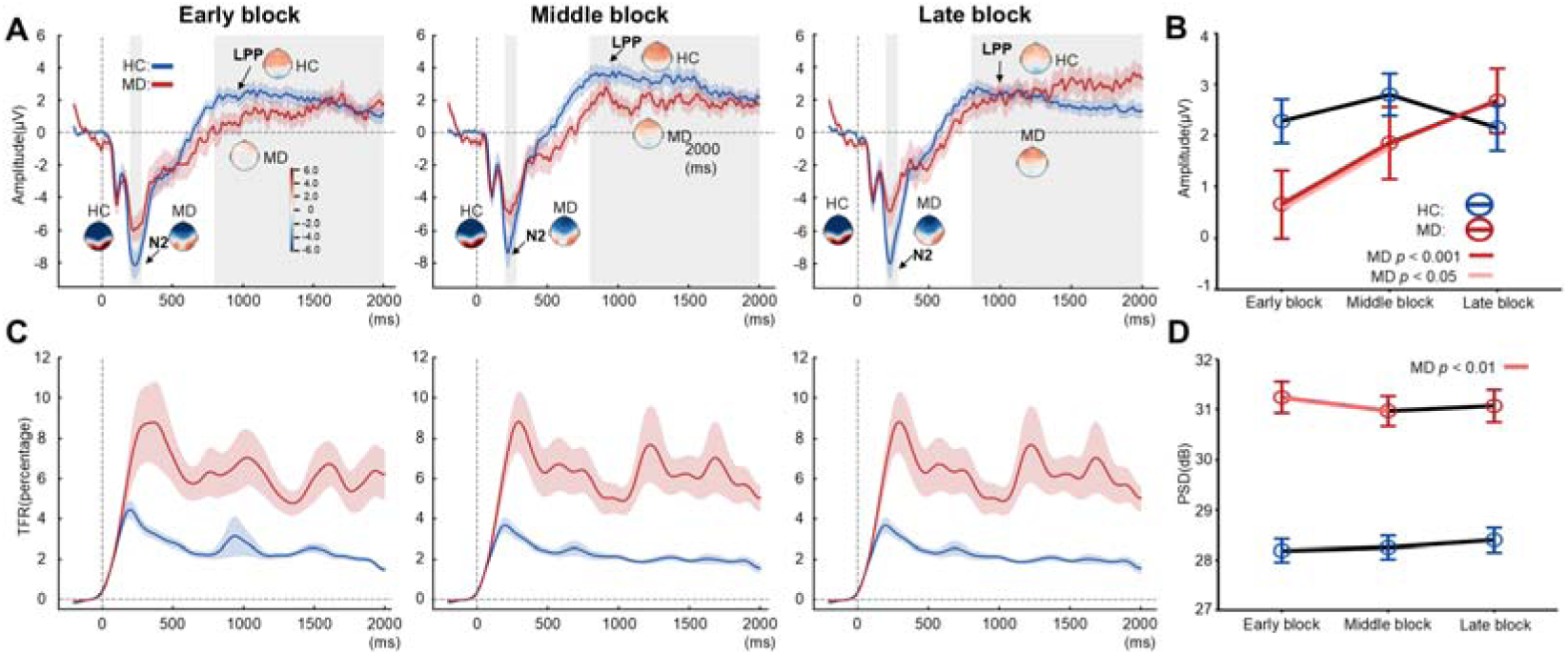
Macro-scale dysregulation: Cumulative emotional sensitization and regulatory depletion. **(A-B) Temporal evolution of the Late Positive Potential (LPP). (A) Grand average ERP waveforms across Early, Middle, and Late experimental blocks. (B) Line chart quantifying mean LPP amplitudes.** Major depression (MD) patients (red) demonstrate a progressive escalation in LPP amplitude as the session proceeds, indicative of cumulative emotional sensitization, whereas Healthy Controls (HC; blue) maintain stable responses. (C–D) Trajectory of frontal Theta activity. **(C) Time-Frequency Representations (TFR) of frontal theta power across the three blocks. (D) Frontal Theta Power Spectral Density (PSD) trajectory.** MD patients exhibit a significant, continuous decline in theta power from early to late blocks, reflecting the cumulative depletion of top-down regulatory resources under sustained exposure. HCs preserve stable regulatory capacity throughout the task. Data are presented as marginal estimated means ± SD. Significance markers indicate group differences: *** *p* < 0.001, ** *p* < 0.01, * *p* < 0.05.

Traveling-wave analysis provided cross-regional dynamic evidence for this micro-scale dysregulation in MD patients (**SI Text 6; Figure 4A**). During the second viewing, HCs exhibited a significant reduction in FW alpha-band propagation velocity (*p* = 0.029, Cohen’s *d* = –0.21). This was interpreted as a typical shift in neural processing from broad, broadcast-like signaling to more refined local processing (Alamia & VanRullen, 2019). MD patients, however, showed no such reduction in the alpha-band propagation velocity.

**Figure 4.**
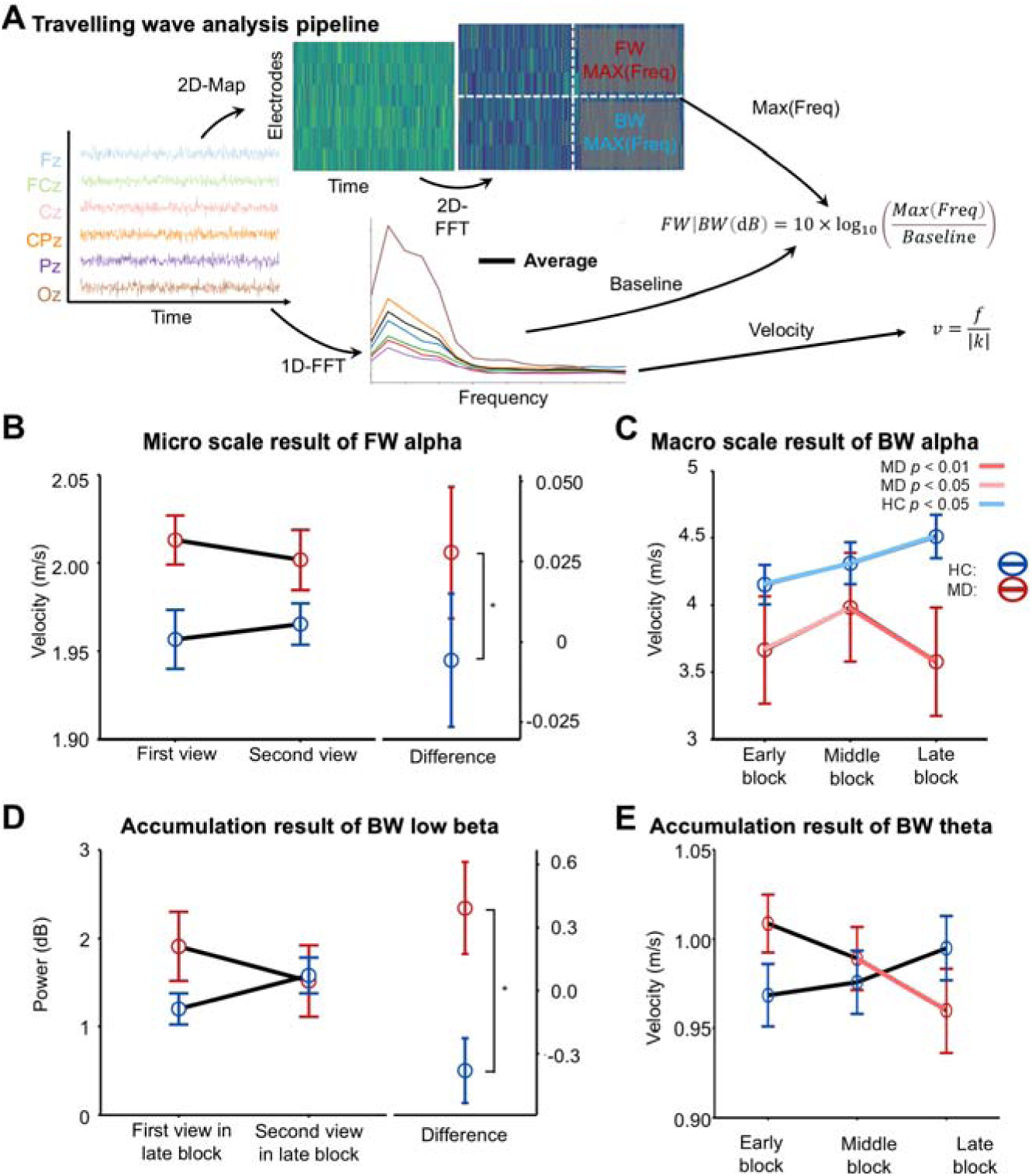
Spatiotemporal dynamics: Multi-scale abnormalities in cortical traveling waves. **(A) Schematic of the Traveling Wave analysis pipeline.** EEG signals from midline electrodes were transformed via 2D-FFT to quantify Forward Waves (FW) and Backward Waves (BW) in terms of power (dB) and phase velocity. **(B) Micro-scale dynamics (FW Alpha Velocity):** Healthy controls (HC) exhibit adaptive slowing of FW Alpha velocity upon repetition (difference < 0), a signature of predictive stabilization. Major depression (MD) patients fail to show this modulation. **(C) Macro-scale dynamics (BW Alpha Velocity):** MD patients show a specific reduction in BW Alpha velocity (not power) during late blocks, indicating a progressive slowdown of feedback signaling over time. **(D**– **E) Cross-scale cumulative interactions. (D)** MD patients develop a paradoxical “runaway” increase in BW Low-Beta Power specifically during the second view of late trials. **(E)** MD patients exhibit progressive deceleration of BW Theta Velocity across the session (Late < Early), reflecting the gradual collapse of top-down regulatory integration. ** *p* < 0.01, * *p* < 0.05.

Overall, these findings converged on a picture of neural rigidity at the micro timescale in MD patients: in the face of repeated stimuli over only a few seconds, rather than rapidly scaling down the neural resources as observed in HCs, MD patients maintained or even reinforced the original response pattern, indicating a fundamental impairment in fast adaptive mechanisms.

#### 2.1.3 Macro-scale cumulative dysregulation over prolonged exposure

Across the full course of the experiment with early-middle-late block, the neural trajectories of HCs and MD patients diverged considerably. Healthy individuals maintained a relatively steady neural response throughout the task, whereas MD patients showed a progressively accumulating trend of disorder in emotional processing. At the ERP level (**SI Text 2; Figure 3A-B**), MD patients displayed an increasing rise in frontal LPP amplitudes across blocks (early < middle < late; Middle-Early: *p_FDR_*= 0.01, *Cohen’s d* = 0.25; Late-Early: *p_FDR_* < 0.001, *Cohen’s d* = 0.42), a trend absent in HCs. This progressive reactivity reflected a typical emotional sensitization process, indicating that instead of becoming less responsive, MD patients grew increasingly reactive to continuous negative stimulation.

At the same time, in parallel with the increase in frontal LPP, theta power exhibited a continuous decrease in both power spectral density (PSD; Middle-Early: *p_FDR_*= 0.036, *Cohen’s d* = −0.33; **SI Text 3; Figure 3C-D**). Because theta rhythm was widely considered a crucial neural index of emotional regulation and cognitive control resources (Cavanagh & Frank, 2014), this downward trend of θ in MD patients indicated that their regulatory systems had shown significant resource depletion under long-term exposure. In contrast, theta activity in HCs remained stable over time, showing no obvious resource depletion in the face of repeated negative stimuli.

Notably, the traveling-wave propagation results provided large-scale network-level evidence for this cumulative dysregulation at the macro timescale (**SI Text 6; Figure 4C**). In HCs, BW alpha energy demonstrated a slight increase in the late block (Late-Early: *p_FDR_* = 0.030, *Cohen’s d* = 0.32), which was consistent with the normal mode of cross-regional information integration that remained stable or even slightly enhanced throughout the task. MD patients, however, showed a marked reduction in BW alpha propagation during the late block (Late-Middle: *p_FDR_* = 0.005, *Cohen’s d* = −0.36), suggesting that feedback signals from higher-order cortical regions to frontal areas became gradually weakened. This pattern indicated a systematic degradation of large-scale networks caused by prolonged exposure to negative stimuli (Greicius et al., 2007; R. H. Kaiser et al., 2015; Muller et al., 2018).

At the macro timescale, our overall results suggested that during long-term exposure to negative stimuli, MD patients not only failed to adapt but also exhibited a worsening trajectory marked by increasingly heightened emotional reactivity, declining regulatory resources, and weakening cross-regional information transmission, a pattern highly consistent with the established models of emotional regulation impairment in MD.

#### 2.1.4 The accumulation of micro-scale deficits to macro-scale dysregulation

To understand how the above micro- and macro-level dysregulations are related, we further examined the three-way interaction among group, time, and block to determine whether patients with MD show a systematic accumulation from micro deficits to macro-scale abnormalities. The results of low beta band energy provided the key evidence (**SI Text 5; Figure 4D**): In the early block, the difference between the first and second view was not significant between MD patients and HCs; however, as the task progressed, MD patients exhibited an abnormal second-view beta runaway in the late block (*p* = 0.047, *Cohen’s d* = −0.33), whereas HCs showed the typical repetition suppression. This three-way interaction revealed that after prolonged exposure to negative stimuli, MD patients not only failed to suppress the neural response to the second presentation, but also showed an overreaction, indicating that their failure of rapid adaptation was continuously amplified at the macro scale (Rubino et al., 2006).

Similarly, the cross-regional propagation dynamics also showed a continuous deterioration from micro to macro levels. BW theta velocity (**SI Text 5; Figure 4E**) did not differ significantly between groups in the early block. However, as the epochs progressed, MD patients showed a distinct gradual slowdown in velocity (*p* = 0.003, Cohen’s *d* = –0.65), while HCs remained stable. This tendency represented a gradual breakdown of the bottom-up pathways involved in emotional integration (Kveraga et al., 2007; Muller et al., 2018). This was considered as an imbalance and overactivation of inhibitory top-down signals (Kveraga et al., 2007), further supporting the notion of a propagation system dysregulation.

Together, the three-way interaction and the diffusion kinetics pointed to a unified picture. Adaptation failure in MD patients was not an isolated occurrence but a deterioration process that spanned across time scales. Micro-scale deficits in rapid suppression during prolonged exposure developed into macro-scale dysregulation in the maintenance of network stability, ultimately forming a neural dynamic trajectory marked by emotional sensitization, regulatory fatigue, and imbalance of cross-regional information flow.

### 2.2 Reverse machine-learning validation

Since multi-scale damage is a more core feature of MD patients, it must contain more damage information than the traditional single shot, so as to achieve better classification performance.

Therefore, at the stage, besides single-shot results, we had four progressively integrated feature sets, which were related to single-shot, micro-scale adaptation, macro-scale adaptation and combined multiscale adaptation. For each feature set, we trained a logistic-regression classifier to distinguish MD patients from HCs using leave-one-out cross-validation (LOOCV). Model performance was assessed with permutation testing, using 100 label-shuffle iterations to generate a null distribution of AUC values (**Table 1**, **Figure 5B-C**).

**Figure 5.**
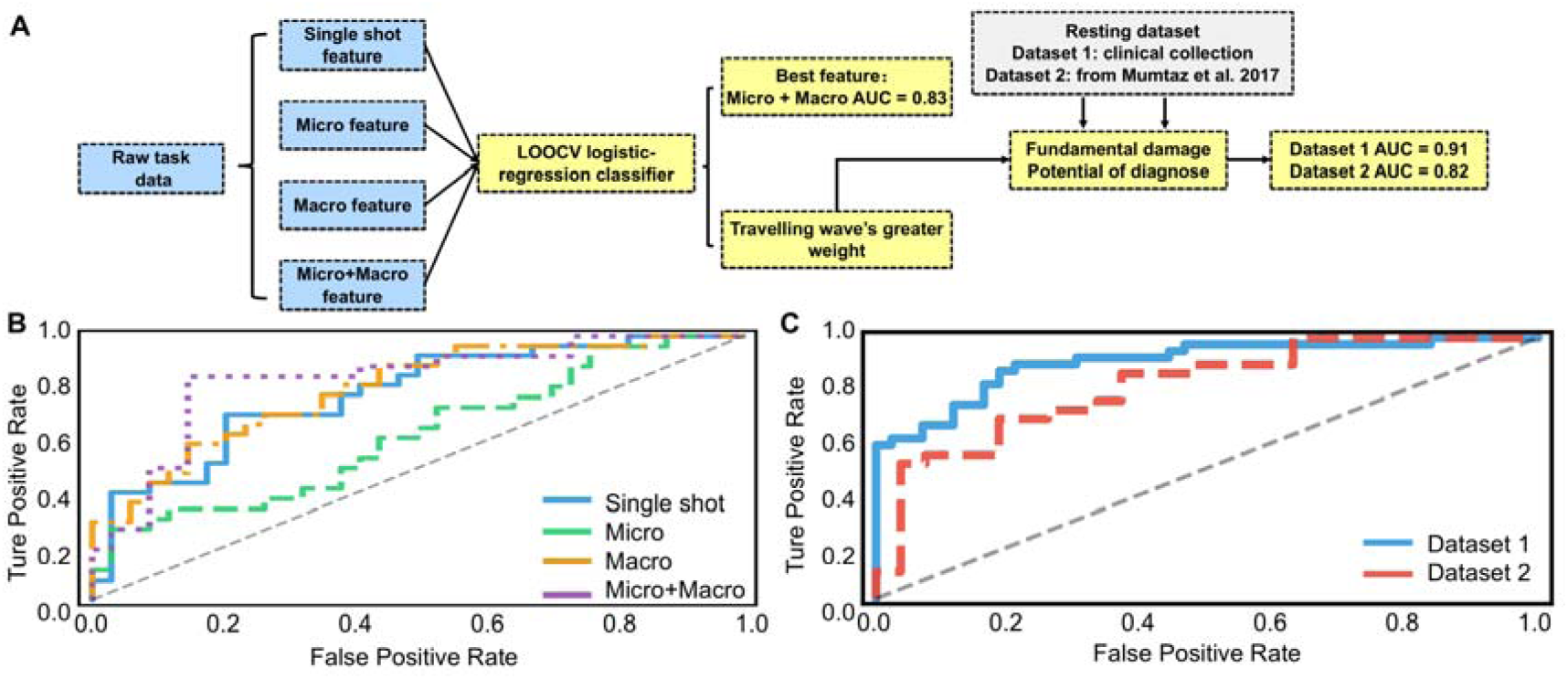
Adaptation-based machine learning classification and cross-dataset generalization. **(A) Machine learning pipeline. Logistic regression classifiers were trained using Leave-One-Out Cross-Validation (LOOCV) on feature sets representing different temporal scales.** The diagram highlights that the combined model yields the best performance (AUC = 0.83) and that traveling wave features contributed the greatest weight to the classification. **(B) Receiver Operating Characteristic (ROC) curves for the task dataset.** The model integrating micro- and macro-adaptation features (purple dotted line) achieved the highest diagnostic performance, significantly outperforming the traditional single-shot baseline (blue solid line; AUC = 0.78). **(C) Out-of-sample generalization to Resting-State.** The traveling-wave adaptation signatures identified in the task robustly generalized to two independent resting-state datasets (Dataset 1: Clinical cohort, AUC = 0.91; Dataset 2: Independent public dataset, AUC = 0.82). This suggests that the identified propagation abnormalities represent a stable, trait-like biomarker of depression distinct from transient task demands.

**Table 1.** Classification performance metrics for four task-state feature sets and two independent resting-state datasets.

| Model | Single shot | Micro | Macro | Micro+Macro | Rest dataset1 | Rest dataset2 |
| --- | --- | --- | --- | --- | --- | --- |
| Accuracy | 0.67 | 0.63 | 0.74 | 0.77 | 0.79 | 0.71 |
| AUC | 0.78* | 0.62 | 0.80* | 0.83* | 0.91*** | 0.82*** |
| Recall | 0.48 | 0.31 | 0.59 | 0.65 | 0.74 | 0.72 |
| Precision | 0.68 | 0.67 | 0.76 | 0.77 | 0.81 | 0.72 |
| F1 | 0.57 | 0.42 | 0.67 | 0.71 | 0.77 | 0.72 |

Task-state models: Single-shot (baseline neural responses), Micro (within-trial adaptation), Macro (temporal habituation trends), and Micro+Macro (combined adaptation features). All models evaluated using leave-one-out cross-validation (LOOCV) with logistic regression. Resting-state datasets serve as independent validation cohorts. Statistical significance determined by permutation testing with 100 iterations. \* $p < 0.05$ ,
\*\*\* $p < 0.001$ .

#### 2.2.1 Hierarchical performance across temporal scales

A clear diagnostic gradient confirmed that MD is fundamentally characterized by **multi-scale adaptation failure** rather than static reactivity anomalies (**Table 1**, **Figure 5B**). While traditional single-shot features provided a moderate baseline (AUC = 0.78, *p* = 0.010), they were outperformed by macro-scale adaptation trajectories (AUC = 0.80, *p* = 0.020), which successfully captured the cumulative depletion of neural resources.

Critically, diagnostic performance peaked only when micro- and macro-scale dynamics were integrated (AUC = 0.83, *p* = 0.010). The fact that the combined model surpassed both static baselines and isolated adaptation scales indicates that the core neural impairment in MD is a compound dysfunction, spanning from rapid micro-scale rigidity to sustained macro-scale regulatory collapse.

#### 2.2.2 Traveling wave dynamics as core diagnostic markers

After identifying the critical role of the multiscale model in MD patient classification, we investigated which types of features drove the outstanding performance of this model. We conducted feature importance analyses and found that traveling-wave parameters contributed more strongly to the classification than traditional ERP measures, accounting for 59.0% of the total coefficient magnitude (cumulative coefficient: 1.10) compared with 41.0% for ERPs (cumulative coefficient: 0.77). This meant that traveling-wave features carried roughly 1.4 times the weight of ERP features in the model, highlighting traveling waves as key carriers of diagnostic information in our multiscale adaptation-based framework.

Within the traveling-wave features, we observed a clear directional asymmetry. Specifically, BW carried substantially larger weight than FW (sum of absolute coefficients: 0.84 vs. 0.27). Also, speed-based metrics outweighed power-based metrics (0.76 vs. 0.35), suggesting that the propagation velocity was a more informative marker for classification than oscillatory amplitude. A frequency-specific analysis further showed that theta-band waves contributed most strongly to classification (0.49), followed by alpha (0.35) and low-beta bands (0.27). This is consistent with the results of our forward analysis. Damage to the low-frequency brain connection network mainly composed of the theta and alpha bands may be the main characteristic of MD patients (Fingelkurts et al., 2007; Sun et al., 2019). This large-scale propagation deficit converged with functional magnetic resonance imaging evidence of decreased functional connectivity in MD patients (R. H. Kaiser et al., 2015).

#### 2.2.3 Generalization to resting-state networks

For the last step, we examined whether the abnormalities of adaptation-related traveling waves in MD patients depended on task-evoked processing or instead reflected more intrinsic properties of neural dynamics. In parallel, because our preceding analysis suggested that traveling-wave features was a more fundamental deficit, we also asked whether these waves could serve as a promising diagnostic biomarker that extended beyond this specific adaptation paradigm. To address these questions, we applied the best-performing adaptation-based model to two independent resting-state EEG datasets with the same feature extraction and classification procedures.

Even though the resting-state recordings did not involve any explicit task, we found traveling-wave adaptation features retained high predictive accuracy (Dataset 1: AUC = 0.91, *p* < 0.001; Dataset 2: AUC = 0.82, *p* < 0.001; **Table 1**, **Figure 5C**). Because the model was trained on task-evoked adaptation features, its consistent performance on resting-state data indicated that the underlying traveling-wave abnormality was task-independent and reflected a stable disturbance in large-scale propagation dynamics in MD patients. Thus, the ability to detect this disturbance from resting-state EEG offered a promising basis for task-free electrophysiological biomarkers.

## 3 Discussion

### 3.1 From momentary abnormalities to adaptation failure: Rethinking the core pathology of MD

Traditional models conceptualize MD primarily through moment-level abnormalities, such as heightened reactivity to aversive information (“negative bias”; Gotlib & Joormann, 2010) or dampened emotional responsiveness (“emotional blunting”; Christensen et al., 2022). Yet these accounts, and the paradigms that generated them, are grounded almost exclusively in single-shot designs that isolate the very first response to a stimulus. These paradigms can reveal *how strongly* a patient reacts at the onset of a negative event, but they cannot reveal *how this response evolves* when the same adversity is repeatedly encountered—an essential feature of real-world depressive experience.

The present findings extend this static view, showing that MD is not defined simply by the magnitude of the first reaction, but by a broader failure of adaptive protection—a pervasive inability to recalibrate neural responses across time. Our data reveal two complementary forms of maladaptation: neural rigidity at the micro scale (2–6 s; absent repetition suppression), and cumulative regulatory depletion at the macro scale (20–30 min; escalating LPP and declining theta). This dual-scale profile provides a mechanistic explanation for why depressive affect remains persistent even when negative events become familiar.

This perspective may offer a reinterpretation of these momentary abnormalities (negative bias or emotional blunting) that characterized MD. Rather than a standalone symptom, momentary abnormalities may represent only the first frame of a longer maladaptive trajectory—the “opening snapshot” of an adaptation process that fails to unfold. Just as a single photograph can capture the start of a movement while obscuring its direction, a single-shot ERP amplitude captures the *initial peak* but not the *trajectory* that follows. In healthy adults, emotional responses typically attenuate through regulatory and predictive mechanisms (Barthel et al., 2025; McEwen, 2000); in MD, our data show they remain elevated or escalate. Thus, momentary abnormalities (negative bias or emotional blunting)—traditionally treated as a defining feature of MD—may instead be the visible tip of a deeper adaptation-failure process, a hypothesis that could not be tested without a paradigm explicitly designed to track neural dynamics across repetitions. This reinterpretation situates MD within a unified plasticity framework: when experience-dependent plasticity fails to recalibrate responses, momentary reactivity like negative bias persists and accumulates, manifesting clinically as sustained negative affect and rumination.

### 3.2 Multi-scale neural signatures of adaptive protection failurew

Our findings reveal a dual-scale maladaptation profile in MD that links short-term repetition suppression failures with long-term deficits in regulatory dynamics.

#### Micro-scale: neural rigidity and failure of repetition suppression

HCs showed clear repetition suppression. LPP amplitude decreased upon re-exposure, and theta power reduced, consistent with efficient disengagement and resource reallocation (Garrido, Kilner, Kiebel, et al., 2009, 2009). In contrast, MD patients showed little to no attenuation and often displayed response potentiation, revealing rigidity in early adaptive circuits. MVPA further corroborated this rigidity. Although classifiers in both groups reliably discriminated first from second viewings, the direction of decoding diverged sharply: in HCs, Haufe-transformed weights were predominantly negative, consistent with an overall reduction of neural activation during the second viewing, whereas in MD patients’ weights were predominantly positive, indicating potentiation of neural responses with repetition rather than suppression. Mechanistically, absent micro-scale attenuation is consistent with reduced inhibitory synaptic reweighting and impaired spike-timing-dependent plasticity windows that normally track stimulus predictability (often theta-gated), preventing efficient disengagement (Turrigiano, 2012; Zenke et al., 2017).

#### Macro-scale: cumulative emotional reactivity and regulatory depletion

Across the whole 20–30 min task session, MD patients showed progressively increasing LPP amplitude and declining theta rhythm energy, consistent with escalating emotional arousal (Cuthbert et al., 2000; Schupp et al., 2000) and diminishing cognitive-regulatory resources (Ang et al., 2023). Crucially, travelling-wave analyses illuminated the network-level mechanism behind these patterns: HCs demonstrated adaptive slowing of FW (alpha), reflecting predictive stabilization (Bahramisharif et al., 2013; M. Halgren et al., 2019), whereas MD patients failed to exhibit such slowing and additionally showed reduced BW (theta) velocity, indicating impaired top-down integration and feedback regulation (Alamia & VanRullen, 2019; A. S. Halgren et al., 2023).

Together, these complementary micro- and macro-scale abnormalities form a coherent pattern: rigid short-term responses that fail to habituate, and long-term trajectories that drift toward exhaustion rather than stability. This integrated profile bridges ERP/oscillation findings with large-scale propagation dynamics, providing the most detailed characterization to date of how adaptation fails in MD over time.

### 3.3 Adaptation-based indices outperform single-shot markers: evidence for mechanistic primacy

If multi-timescale adaptation failure is indeed the mechanistic bottleneck of MD, then adaptation-based measures should necessarily outperform single-shot indices—both in explaining larger group differences (forward comparison) and in yielding superior diagnostic classification (reverse comparison). Our data confirmed this prediction across levels of analysis. For the forward comparison, micro-scale adaptation slopes (changes in LPP and theta activity) showed larger and clearer group differences than initial amplitudes (replicating but exceeding the static deficits reported in Jin et al., 2022; Stone et al., 2025), and macro-scale trajectories (LPP increases and theta declines across the session) likewise revealed robust separations that single-shot responses could not detect. Traveling-wave adaptation metrics, including changes in FW and BW velocity, produced some of the strongest group contrasts, indicating that large-scale propagation dynamics capture discriminative information that momentary responses fail to reveal (Alamia et al., 2024; Alamia & VanRullen, 2019).

The reverse comparison yielded a parallel pattern: adaptation-based classifiers markedly outperformed single-shot classifiers. In the task dataset, the adaptation model achieved an AUC of 0.83, compared with 0.78 for models relying solely on single-shot indices (a performance ceiling often observed in static feature benchmarks; e.g., Kaushik et al., 2023). This advantage generalized to an independent clinical replication dataset, where the adaptation model reached an AUC of 0.91, and to a public resting-state dataset (Mumtaz et al., 2017), where the adaptation model achieved an AUC of 0.83. Across datasets, traveling-wave components—particularly BW velocity—consistently ranked among the most informative features, underscoring their reproducibility and robustness as biomarkers.

Taken together, these findings demonstrate that adaptation captures richer and more diagnostic information than traditional single-shot measures. This supports the theoretical interpretation developed earlier: what appears as negative bias in isolated trials may be only a visible slice of a deeper adaptation deficit, whereas adaptation-based indices capture the temporal dynamics through which MD pathology more fundamentally unfolds.

### 3.4 Mechanistic and theoretical implications: From rumination and cognitive inflexibility to predictive dysregulation

The dual□scale maladaptation pattern identified here—micro□scale neural rigidity and macro□scale regulatory depletion—may offer a mechanistic bridge to hallmark symptoms of MD. At short timescales, impaired repetition suppression (sustained LPP and theta activity) provides a plausible neural substrate for why repetitive negative thoughts fail to habituate, contributing to *rumination* (Gotlib & Joormann, 2010; Zhou et al., 2020). At longer timescales, the progressive escalation of LPP amplitude coupled with declining theta power aligns with well-documented deficits in cognitive flexibility, including difficulty shifting attention or updating emotion regulation strategies under prolonged stress (Aldao et al., 2015; Hu & Tamir, 2025; Murphy et al., 2012). This two-stage pattern—early rigidity cascading into later exhaustion—illustrates how momentary persistence of neural reactivity can accumulate into enduring psychological entrapment.

Beyond symptom-level interpretation, these findings integrate naturally with predictive coding and dynamical systems frameworks. Within predictive coding, repetition suppression reflects decreasing prediction error as stimuli become expected (Friston, 2018). The absence of suppression in MD, alongside aberrant striatal prediction-error signaling reported previously (Gradin et al., 2011; Kumar et al., 2008), suggests impaired hierarchical updating or dysfunctional precision weighting. Traveling waves further illuminate the spatial implementation of these temporal dynamics: healthy individuals showed adaptive slowing of FW (alpha) propagation—consistent with stabilization of predictive templates (Alamia & VanRullen, 2019; M. Halgren et al., 2019)—whereas MD patients displayed persistent forward velocity and reduced backward (theta) propagation, indicative of weakened feedback integration. Within hierarchical predictive coding, such a pattern implies faulty precision-weighting and feedback-driven synaptic plasticity: alpha-/theta-band backward waves normally convey predictions and top-down gain control; their slowing/weakening indicates diminished feedback plasticity across cortical hierarchies (Friston, 2018; Halgren et al., 2019).

### 3.5 Clinical implications and future directions

The present findings carry several conceptual and translational implications. By demonstrating that MD pathology is characterized not simply by heightened initial reactivity but by a pervasive failure of adaptive protection across time, our results point toward new therapeutic targets that extend beyond modulating momentary emotional responses. Interventions that reduce the frequency or duration of repetitive negative exposure—such as behavioral activation (Dimidjian et al., 2006; Jacobson et al., 1996)—may help mitigate the cumulative emotional burden that our macro-scale results reveal. Likewise, treatments that enhance neural plasticity, including transcranial magnetic stimulation or cognitive control training (George et al., 2010; Hoorelbeke & Koster, 2017), may directly address the micro-scale rigidity observed in early neural responses.

From a practical standpoint, traveling wave analysis offers substantial clinical advantages. Reliable measurements can be obtained with a minimal electrode setup (No more than eight midline electrodes), significantly reducing implementation complexity. Furthermore, traveling wave uses stable electrodes and need no time windows choosing, therefore, the analytic pipeline for traveling wave parameters is more standardized, compared to traditional ERP analyses, which typically rely heavily on expert knowledge and experience (Clayson et al., 2025). This standardization decreases analytical complexity, enabling easier learning and facilitating mass-scale analysis. Consequently, traveling wave analysis has the potential to become a low-cost yet high-performance diagnostic biomarker, particularly valuable in large-scale screening initiatives in resource-limited settings, addressing a crucial global need for accessible mental health diagnostic tools (Hodgkinson et al., 2017).

### 3.6 Conclusion

The present study identifies pervasive neural maladaptation as a defining feature of MD across both immediate and prolonged time scales. Rather than exhibiting the expected attenuation with repeated exposure, MD patients showed rigid, persistently heightened responses at the micro scale and progressive emotional escalation with regulatory depletion at the macro scale. These temporal abnormalities were robustly detected by adaptation-based neural indices—particularly traveling-wave dynamics—which outperformed conventional single-shot measures and generalized reliably across independent datasets.

Conceptually, these findings shift the understanding of MD pathology away from static accounts centered on initial hyper-reactivity. Instead, MD emerges as a disorder of impaired temporal adaptation, in which emotional responses fail to recalibrate despite repetition or familiarity. Within this framework, the widely discussed “negative bias” can be understood not as a standalone feature, but as the early expression of a maladaptive trajectory that remains uncorrected over time. This perspective unifies momentary neural responses with their temporal evolution and illuminates a mechanistic pathway through which persistent negative affect, rumination, and cognitive rigidity may arise.

## 4 Methods

### 4.1 Participants

MD patients were recruited from a psychiatric hospital [name to be disclosed upon acceptance]. All patients were diagnosed by clinicians using the Structured Clinical Interview for DSM Disorders (SCID; First, 2002) was diagnosed as a MD episode, according to the DSM-5 (American Psychiatric Association, 2022). The exclusion criteria for patients in this study included: (1) current or previous neurological disease; (2) any comorbidities of Axis I mental disorders; (3) history of epilepsy or traumatic brain injury. Participants in the healthy control (HC) group were recruited through online advertisement, matched on sex and age. All participants were right-handed and had normal or corrected-to-normal vision. The experimental protocol was approved by the Ethics Committee of the hospital (Approval number: 2020-38-department) and strictly followed the ethical requirements of the Declaration of Helsinki and its subsequent revisions. All MD patients and HCs signed written informed consent prior to the start of the experiment and received cash payment or free psychological counseling services after the experiment.

For task data, after excluding participants who withdrew during the study and those with excessive head movements that significantly affected the quality of the EEG data, a total of 28 MD patients (7 males, 21 females, mean age = 25.93 ± 5.71) and 34 HCs (13 males, 21 females, mean age = 23.09 ± 6.10) were included in the further analysis. G*Power (version 3.1.9.7; with the following parameters: α = 0.05, *power* = 0.80, effect size *f* = 0.25) revealed that a minimum of 56 subjects would be required to detect a moderate effect size (*Cohen’s f* = 0.25). Therefore, the sample size (a total of 62 participants) in this study is sufficient.

For rest data, we recruited 40 MD patients (12 males, 28 females, mean age = 26.98 ± 5.39) and 42 HCs (17 males, 25 females, mean age = 23.98 ± 6.78). All demographic information is shown in **Table 2**.

**Table 2.** Demographic information.

| Item | HC ($M \pm SD$) | MD ($M \pm SD$) | $\chi^2 / t$ |
| --- | --- | --- | --- |
| **Task data** |  |  |  |
| Gender(male/female) | 13/21 | 7/21 | 0.70 |
| Age | 23.09 ± 6.10 | 25.93 ± 5.71 | -1.88 |
| BDI | 4.88 ± 4.82 | 22.04 ± 11.91 | -7.68*** |
| **Rest data** |  |  |  |
| Gender | 17/25 | 12/28 | 0.73 |
| Age | 23.98 ± 6.78 | 26.08 ± 5.39 | -1.51 |
| BDI | 5.37 ± 4.68 | 22.04 ± 11.91 | -14.39*** |

BDI: Beck-Depression Inventory. \*\*\*: $p < 0.001$

To further test generalizability, we additionally analyzed an independent public EEG dataset from Mumtaz et al. (2017). This dataset contains resting-state EEG recordings from 34 patients diagnosed with MD and 30 age-matched HCs, acquired using a standard 19-channel international 10–20 montage. All participants were confirmed by structured clinical interviews following DSM criteria. We used this dataset exclusively for out-of-sample validation of the key propagation markers identified in our main experiment.

### 4.2 Experimental design

The experimental design is illustrated in **Figure 1A**. In each trial, participants were required to view the same negative stimulus twice. Each trial began with a fixation cross presented for 600–800 ms, followed by the first view task lasting 5000 ms. Afterward, there was a break with an empty screen for 2000 ms. The second view task, involving the same stimulus, was then presented for another 5000 ms. Finally, participants provided a negative rating (on a scale of 1–9, with higher scores indicating more negative emotions perceived). Each participant completed a total of 120 trials. All visual stimuli were validated negative images from the International Affective Picture System (IAPS; Lang, 1995), preselected based on normative ratings (*M*_valence_ = 2.84 ± 0.77; *M*_arousal_ = 5.46 ± 0.88).

### 4.3 Behavioral data

We recorded the participants’ emotional ratings (on a scale of 1–9, with higher scores indicating more negative emotions perceived) at the end of each trial for statistical analysis.

### 4.4 EEG data recording and processing

EEG data were recorded during the view block and the distraction block using a 32-channel amplifier (Brain Products, Gilching, Germany), with a sampling frequency of 250 Hz. Electrode impedances were kept below 10 kΩ. The reference electrode was placed at the FCz.

All EEG data were preprocessed in Python using the MNE library version 1.7.1 (Larson et al., 2024). Bad channels were first manually inspected and interpolated, then data were averaged to common average and filtered using a 0.1 to 30 Hz bandpass filter (Butterworth order 3) and a 50 Hz notch filter. The data were segmented into epochs of −200-2000ms. To correct eyeblink artifacts, data were processed through MNE’s independent component analysis (ICA) without interpolated channels. Components with blink artifacts were automatically detected by MNE-ICALabel toolbox (Li et al., 2022). Then we used autoreject version 0.4.2, a Python-based data-driven method that learns the maximum peak-to-peak thresholds based on trials from each subject to interpolate the bad channels and reject epochs (Jas et al., 2017). Finally, we corrected the data within a window from −200 ms to 0 ms. All automated results were manually reviewed. We visually examined the distribution of retained epochs per subject to ensure that each subject had at least 20 retained epochs per trial parts.

### 4.5 Temporal neural dynamics

#### 4.5.1 Event-related potentials (ERPs)

Event-related potentials (ERPs) were identified as the mean amplitude in specific time windows, guided by previous literature and adjusted based on visual inspection of the grand average waveforms. N2 was extracted from the frontal region (F3, F4, Fz) between 200-280 ms (Folstein & Van Petten, 2008) and LPP components were calculated in the 800-2000 ms respectively, using the frontal region electrodes (F3, F4, Fz; Hajcak et al., 2010). Frontal LPP has been confirmed to have a direct correlation with view effect (Dennis & Hajcak, 2009; Hajcak et al., 2010), especially in MD patients (Waters & Tucker, 2016). Frontal ERPs in N2 and LPP time windows were averaged for statistical analysis.

#### 4.5.2 Multivariate pattern analysis (MVPA)

To overcome electrode selection bias inherent in traditional ERP analyses that rely on limited electrode sites (Grootswagers et al., 2017), we employed multivariate pattern analysis (MVPA). This approach characterizes neural activity patterns across the entire electrode array, providing a more comprehensive measurement of the repetition suppression (Carrasco et al., 2024). MVPA on the preprocessed EEG data was applied using the MNE and scikit-learn (Larson et al., 2024; Pedregosa et al., 2012) with a logistic regression classifier. We categorized trials into “first view” vs. “second view” classes and tested whether the classifier could learn to discriminate these two conditions based on distinct EEG patterns. We used five-fold cross-validation within each participant: the dataset was split into five-fold, the classifier was trained on four folds and tested on the remaining one. To further reduce selection bias, the entire process was repeated five times with a new random division each time. When necessary, we down-sampled by randomly discarding trials to ensure equal trial numbers in both conditions (Jing-Hao Xue & Hall, 2015). Classification performance was assessed via the area under the ROC curve (AUC), a measure from signal detection theory that is relatively insensitive to classifier bias (Bradley, 1997). After decoding, one-tailed t-tests against 50% chance level were performed on the group-level AUC at each time point, followed by cluster-based permutation tests (*p* < 0.05, 1000 iterations) to correct for multiple comparisons. Additionally, a temporal generalization analysis was conducted to examine the stability of neural activity patterns across time (King & Dehaene, 2014). The classifier trained at a specific time point was tested on all other time points; above-chance accuracy off the main diagonal indicated stable neural patterns. Lastly, we employed Haufe’s method to convert linear classifier weights into interpretable activation patterns, facilitating further interpretation of the MVPA results (Haufe et al., 2014).

### 4.6 Neural oscillation

#### 4.6.1 Frequency-domain analysis

The theta rhythm is considered a key indicator of the cognitive effort invested by participants during the task (Ang et al., 2023). Previous studies show that theta rhythm is most pronounced in the prefrontal area, a finding confirmed in both humans and animals (Cavanagh & Frank, 2014).

First, to quantify overall oscillatory power, Power Spectral Density (PSD) was estimated using Welch’s modified periodogram method. The EEG time series were segmented into overlapping temporal windows, and a Hanning window was applied to each segment to mitigate spectral leakage. Fast Fourier Transform (FFT) was then computed for each window, and the resulting periodograms were averaged to yield a robust estimate of power distribution. The PSD values were log-transformed to normalize the distribution. For statistical analysis, we averaged the PSD values across the frontal region of interest (F3, F4, Fz) and integrated power within the theta frequency band (4–8 Hz) to derive a single mean power index per condition. Second, to capture dynamic changes within the trial window, we conducted Time-Frequency Analysis using the Morlet wavelet transform. The analysis window was extended (−500 ms to 2300 ms) to minimize edge effects (Iemi et al., 2017). Frequencies were set from 4 to 8 Hz (integer steps), with the number of cycles set to half the frequency value to optimize the trade-off between temporal and spectral resolution. Responses were baseline-corrected (−200 to 0 ms) and expressed as percentage change. This time-resolved approach allowed us to track the trajectory of theta activity throughout the stimulus presentation. We visualized the intra-trial activities through time-frequency analysis (TFR).

#### 4.6.2 Travelling wave energy

The analysis pipeline is illustrated in **Figure. 5A**, adapting from methodologies outlined in (Alamia et al., 2024; Orsher et al., 2024). We quantified traveling wave’s propagation along 6 midline electrodes, running from occipital to frontal regions (Oz, Pz, CPz, Cz, FCz, and Fz). We stacked the signals from the 6 electrodes to create 2D maps, with time and electrodes as axes. From each map, we computed 2D FFT: importantly, the power in the lower and upper quadrants quantifies the amplitude of forward wave (FW; from occipital to frontal electrodes) and backward wave (BW; from frontal to occipital) waves, respectively. FW reflects perception and attention based on visual stimuli (Wilson et al., 2001), and BW reflects top-down adaptive control and adaptive anticipation (Alamia et al., 2024). For each frequency in the 4 to 30 Hz range, we considered the maximum values in both the upper and lower quadrants, obtaining a spectrum for both BW and FW, respectively. We then obtained a baseline value using average results of 1D-FFT and converted FW and BW into dB units based on the baseline value. The formula for calculating traveling wave is defined as follows:

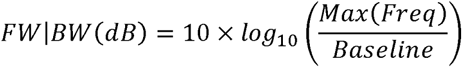

The baseline was the mean of the 1D-FFT results. BW was calculated in the upper quadrant and FW was calculated in the lower quadrant. For the theta (4-8Hz), alpha (8-14Hz), low beta (14-20Hz), and high beta (20-30Hz) rhythms contained in 4-30Hz, we obtained average values respectively for subsequent statistical tests and single trial values for machine learning.

#### 4.6.3 Travelling wave velocity

In addition to spectral power, we quantified the propagation speed of travelling waves. For each subject and condition, multi-channel EEG data from six midline electrodes (Fz, FCz, Cz, CPz, Pz, Oz) were transformed using a 2D-FFT across the spatial and temporal dimensions within a sliding time window. FW and BW components were separated by the sign of the spatial frequency. For each frequency bin, the dominant spatial frequency (k, cycles/m) was identified, and the corresponding phase velocity was computed as

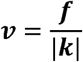

where f is the temporal frequency (Hz). Velocity estimates were then averaged within canonical frequency bands (theta: 4–7 Hz, alpha: 8–13 Hz, low beta: 14–20 Hz, high beta: 21–30 Hz) and across trials.

### 4.7 Statistical analysis and machine-learning validation

Our analytical strategy comprised two complementary steps: a forward statistical inference to compare single-shot versus adaptation metrics, and a reverse machine-learning validation to assess diagnostic utility and generalizability.

#### 4.7.1 Forward statistical inference

All statistical analyses were conducted in R 4.2.2 using the *bruceR* package (Bao, 2024). Simple-effect analyses and multiple comparisons were corrected using the false discovery rate (FDR; (Cao & Zhang, 2014).

##### Single-shot analysis (baseline responses)

To reproduce conventional single-shot approaches, we extracted initial neural responses—the amplitudes/power of N2, LPP, theta rhythms, travelling-wave parameters, and behavior performance—only from the first presentation of each stimulus. These values represent the baseline responding prior to any repetition and thus constitute the single-shot index. Group differences (HC vs. MD) were tested using independent-samples *t*-tests.

##### Micro-scale adaptation (within-trial repetition suppression)

To examine short-timescale adaptation, we quantified changes from first view → second view within each trial. Micro-adaptation indices included: Δ N2 amplitude, Δ LPP amplitude, Δ theta power, and Δ traveling-wave parameters (Δ power, Δ velocity in FW/BW directions)

A mixed-design ANOVA was performed with view (first vs. second) and block (early, middle, late) as within-subject factors and group (HC vs. MD) as a between-subject factor. Marginal estimated means (±1 *SD*) are plotted in line charts.

##### Macro-scale adaptation (across-session habituation trajectory)

Macro-adaptation indices quantified how neural responses changed over the entire 120-trial session, independently of micro-scale repetition. These included: negative emotion scores, LPP habituation slopes (change across blocks), theta power decline slopes, and traveling-wave trajectory slopes (FW/BW power and velocity).

Block-based trajectories were analyzed using mixed-design ANOVA with block (early, middle, late) as the within-subject factor and group as the between-subject factor.

#### 4.7.2 Reverse machine-learning validation

To test whether adaptation features outperform static responses in distinguishing MD from HC, we compared four feature sets: single-shot (initial responses only), micro-adaptation (first→second changes), macro-adaptation (block-level trajectories), and combined adaptation (micro + macro). Each set included ERP/oscillation measures and traveling-wave features appropriate to the adaptation scale (e.g., only slopes for macro-scale).

##### Modeling procedure

Classification was performed using L2-regularized logistic regression with class-balanced weights and leave-one-out cross-validation (LOOCV). Within each fold, the top 5 discriminative features were selected using univariate ANOVA to reduce overfitting. Performance metrics included AUC, accuracy, precision, recall, and F1-score. Statistical significance was assessed using 100-iteration permutation testing; models with *p* < 0.05 were considered above chance.

##### Feature-importance analysis

For the best-performing model, we conducted feature importance analysis by examining standardized logistic regression coefficients. Features were categorized by type (ERP vs. traveling wave) and, for traveling wave features, further stratified by propagation direction (forward vs. backward), metric type (power vs. speed), and frequency band (theta, alpha, beta-low, beta-high) to quantify their relative contributions to classification accuracy.

##### Generalization to independent datasets

To evaluate whether adaptation-relevant propagation signatures reflect task-dependent vs. intrinsic neural dysfunction, the same traveling-wave extraction and classification pipeline was applied to: a self-collected clinical cohort (34 MD, 30 HC), and a public resting-state dataset (Mumtaz et al., 2017). This served as out-of-sample validation to test generalizability.

## Acknowledgments

This work was supported by the Shenzhen Scientific Program (RCBS20231211090516018; 827000853), and Shenzhen-Hong Kong Institute of Brain Science (2023SHIBS003).

## Disclosures

The authors report no financial interests or potential conflicts of interest.

## Data availability

The datasets generated and analyzed during the current study are available in the Open Science Framework (OSF) repository: https://osf.io/dum3x/overview?view_only=2c760ddf4a3247b596cd29ab9f57c1ab. The custom code used for data analysis is available on GitHub: https://github.com/yykaiii915-coder/Maladaptation-EEG-code.git.

---

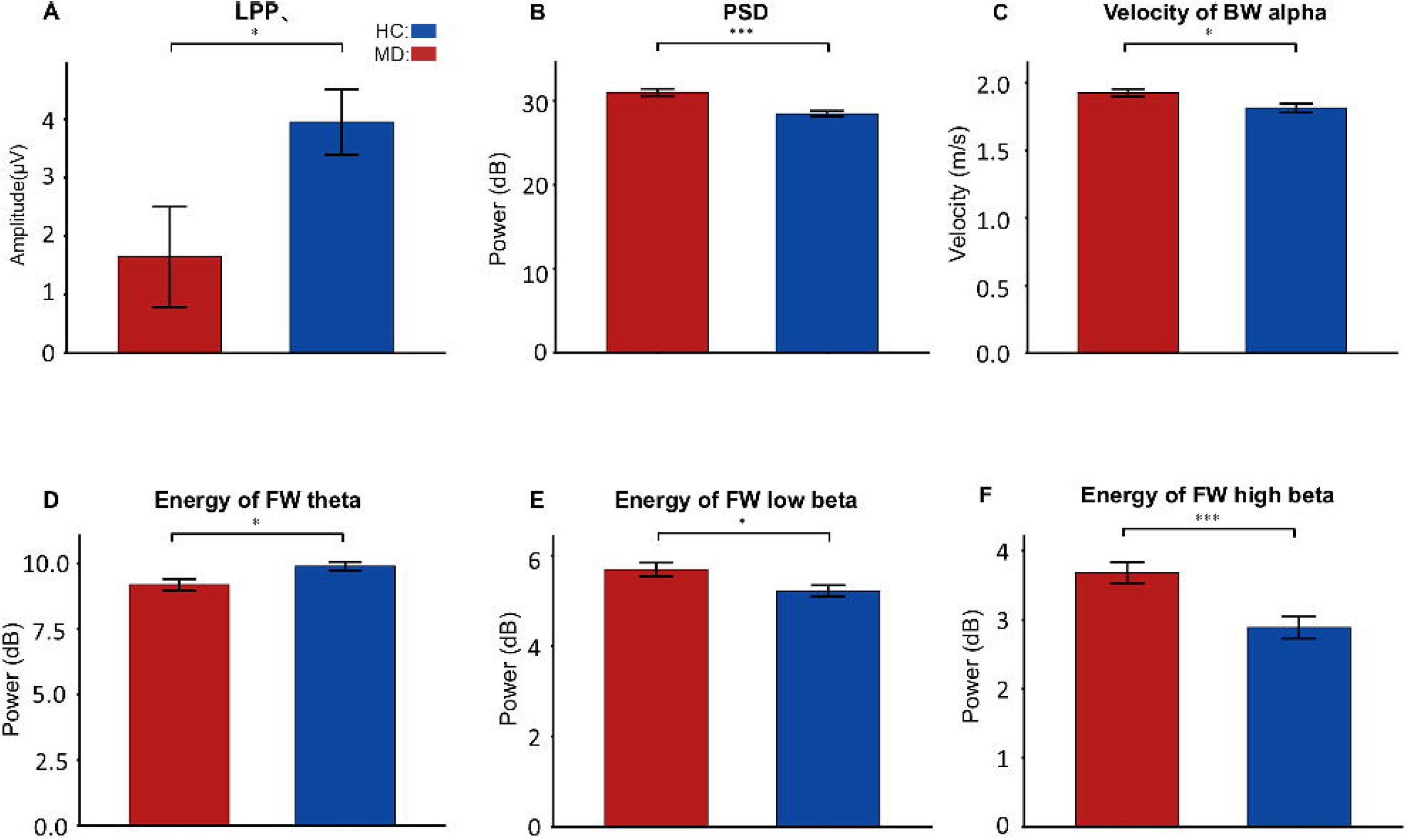

## Notes

### Competing Interest Statement

The authors have declared no competing interest.

