## Supporting Information for "Multi-Timescale Neural Adaptation Failure as a Mechanistic Signature of Major Depression"

Text 1. Behavioral data

We only found the main effect of *group* (*F_(1,60)_* = 5.05, *p* = 0.028, *η^2^p* = 0.08). The negative rating of MD patients was significantly higher than that of HCs (*t*_(60)_ = 2.25, *p* = 0.028, *Cohen’s d* = 1.18). The mean and standard deviation of the negative rating in each condition are shown in **SI Table S1**.

Text 2. Time-domain analysis

**Single shot:** For the frontal LPP in the single-shot test, we observed a significant difference, indicating that the MD group exhibited a smaller LPP amplitude than the HC group (*t*_(60)_ = -2.25, *p* = 0.029, *Cohen’s d* = -0.59**; Figure S1 A**). Statistical analysis of the N2 revealed no significant difference, with a non-significant trend indicating a larger N2 amplitude in the MD group compared to the HC group (*t*_(60)_ = 0.95, *p* = 0.346, *Cohen’s d* = 0.24).

**Microscale**: For frontal LPP (**SI Table S2**), we observed a main effect of *time* (*F*_(1,60)_ = 7.20, *p* = 0.009, *η^2^p* = 0.11) and significant differences between groups on the microscale. A significant *group***time* interaction was found (*F*_(1,60)_ = 8.87, *p* =0.004, *η^2^p* = 0.13). Specifically, the LPP in HCs significantly decreased during the second view time (*t*_(60)_ = -4.21, *p* < 0.001, *Cohen’s d* = -0.64), whereas no such change was observed in MD patients (*t*_(60)_ = 0.20, *p* = 0.842, *Cohen’s d* = 0.03).

For N2, we found that MD patients and HCs exhibited similar changes across microscale. A significant main effect of *time* (*F*_(1,60)_ = 8.63, *p* = 0.005, *η^2^p* = 0.13) was found. *Post-hoc* comparison showed that N2 was more negative in the first time of view (*t_(60)_* = 2.94, *p* = 0.005, *Cohen’s d* = 0.39). The statistical results and average amplitude of LPP and N2 at the micro scale are illustrated in **Figure 2A and Figure 2B**.

**Macroscale****:** For LPP, we observed a significant difference between groups on the macroscale. A significant *group***part* interaction was found (*F*_(2,120)_ = 5.57, *p* = 0.005, *η*^2^*p* = 0.09). *Post-hoc* comparison revealed that LPP amplitude increased across parts in MD patients (Middle-Early: *t*_(60)_ = 2.67, *p_FDR_* = 0.01, *Cohen’s d* = 0.25; Late-Early: *t_(60)_* = 4.24, *p_FDR_* < 0.001, *Cohen’s d* = 0.42), while remaining relatively stable in HCs (Middle-Early: *t*_(60)_ = 1.28, *p_FDR_* = 0.309, *Cohen’s d* =0.11; Late-Early: *t_(60)_* = -0.30, *p_FDR_* = 0.764, *Cohen’s d* = -0.03). The interaction of *group***part* on LPP is depicted in **Figure 3A**, with the average LPP oscillogram across three parts shown in **Figure 3A**. The mean and standard deviation of the LPP in each condition are shown in **SI Table S2**.

For N2, we found that MD patients and HCs exhibited similar changes across macroscale. A significant main effect of *part* (*F*_(2,120)_ = 5.80, *p* = 0.004, *η^2^p* = 0.09) was found. N2 amplitude was decreased over the part of the experiment (Middle-Early: *t*_(60)_ = 3.42, *p_FDR_* = 0.003, *Cohen’s d* = 0.20; Late-Early: *t*_(60)_ = 2.49, *p_FDR_* = 0.023, *Cohen’s d* = 0.14). **Figure 3A** displays the *group***part* interaction for N2, while **Figure 3A** shows the average N2 oscillogram across three parts. The mean and standard deviation of the N2 in each condition are shown in **SI Table S3**.

Text 3. PSD analysis

**Single shot:** For PSD component in the single-shot test, we observed a significant larger PSD amplitude in the MD group than that in the HC group (*t*_(60)_ = 4.04, *p <* 0.001, *Cohen’s d* = 1.12; **Figure S1 B**).

**Microscale****:** We observed inter-group differences in PSD across microscale. A significant *group***time interaction was found* (*F*_(1,60)_ = 8.39, *p* = 0.005, *η^2^p* = 0.12). While HCs showed a significant reduction in PSD during the second view time (*t*_(60)_ = -2.62, *p* = 0.011, *Cohen’s d* = -0.39), MD *patients exhibited no such reduction. The interaction of group*time* on PSD is illustrated in **Figure 2C-D**.

**Macroscale****:** Significant group differences were also detected on macroscale, with a notable *group***part* interaction (*F*_(2,120)_ = 3.54, *p* = 0.032, *η^2^p* = 0.06). Simple effect analysis revealed that MD patients experienced a decrease in theta rhythm energy (Middle-Early: *t*_(60)_ = -2.59, *p_FDR_* = 0.036, *Cohen’s d* = -0.33), a pattern not observed in HCs (Middle-Early: *t*_(60)_ = 0.67, *p_FDR_* = 0.504, *Cohen’s d* = 0.08). Statistical results for the macro scale are presented in **Figure 3B**. A significant main effect of *group* was found _(1,60)_ = 28.05, *p* < 0.001, *η^2^p* = 0.32), and MD patients have larger theta rhythm energy (*t*_(60)_ = 5.30, *p* < 0.001, *Cohen’s d* = 3.34). The mean and standard error of the PSD in each condition are shown in **SI Table S4**.

**Fatigue check:** We checked the PSD of alpha rhythm at the Cz electrode, which can reflect the fatigue level (Tran et al., 2020). We used a three-way ANOVA with *time* of view (first, second) and *part* of experiment (early, middle, late) as within-subject factors, and *group* (HCs, MD) as a between-subject factor. We found a relatively consistent trend in MD patient and HCs and did not find the interaction of *group***part* (*F*_(2,120)_ = 0.40, *p* = 0.674, *η^2^p* = 0.007). This indicates that there is no difference in the degree of fatigue between MD patients and HCs. This consistent fatigue response cannot be the reason for the differentiation between MD patients and HCs on the macro time scale.

Text 4. EEG MVPA results

MVPA revealed a significant above-chance difference between classes of the first view and second view from 0 to 2000ms (*p* < 0.05, cluster-corrected) both in HCs and MD. Because successful classification was observed in all four data-sets, in the next step, the temporal generalization matrices were calculated to test the stability of neural activity patterns with underlying significant classification performance. The time generalization matrices showed that significant above-chance activity was observed during the significant time windows acquired from the MVPA, suggesting that the differences between pairs of conditions were stable over time. Although MVPA can be successfully decoded in both HCs and MD, the decoding direction is not the same. As shown in the **Figure S1**, decoding weights in HCs are generally negative, while decoding weights in MD are mostly positive. This reflects a reduction in neural activity from the first view to the second view in HCs and the opposite in MD.

Text 5. EEG travelling wave energy results

**Single shot**

**FW energy:** For the energy of FW in theta band, we found significantly smaller amplitude in the MD group than that in the HC group (*t*_(60)_ = -2.62, *p* = 0.012, *Cohen’s d* = -0.68). Similar significant larger amplitudes were also found in low beta band (*t*_(60)_ = 2.42, *p* = 0.019, *Cohen’s d* = 0.63), and to a greater extent in high beta band (*t*_(60)_ = 3.61, *p* < 0.001, *Cohen’s d* = 0.10).

**Energy of BW**

**Macroscale:** For the BW in the alpha band, we found a significant main effect of *part* (*F*_(2,120)_ = 3.73, *p* = 0.027, *η^2^p* = 0.06) and an interaction of *group*part* (*F*_(2,120)_ = 6.48, *p* = 0.002, *η^2^p* = 0.10). Post-hoc test found that the BW energy in the middle part was significantly higher than that in the early part (*t*_(60)_ = 3.10, *p* = 0.009, *Cohen’s d* = 0.21). Simple effect analysis shows the BW energy increased from early part to late part in HCs (Late-Early: *t*_(60)_ = 2.66, *p_FDR_* = 0.030, *Cohen’s d* = 0.32), but in MD, it decreased in the late part (Late-Middle: *t*_(60)_ = -3.29, *p_FDR_* = 0.005, *Cohen’s d* = -0.36). The interaction is shown in the **Figure 4C**.

For the BW in the low beta band, we found a significant interaction of *group*time*part* (*F*_(2,120)_ = 4.51, *p* = 0.013, *η^2^p* = 0.07). Simple effect analysis found that the ES effect of BW in the late part was different in HCs (*t*_(60)_ = 2.16, *p* = 0.035, *Cohen’s d* = 0.32) and MD (*t*_(60)_ = -2.02, *p* = 0.047, *Cohen’s d* = -0.33). For the BW in the high beta band, we did not find any significant main effect (All non-significant effect: *F*_(1,60)_ < 1.13, *p* > 0.292, *η^2^p* < 0.02). The interaction is shown in the **Figure 4D**. The mean and standard deviation of the BW in each condition are shown in **SI Table S9 – S12.**

Text 6. EEG travelling wave velocity results

**Single shot**

**BW velocity:** For the velocity in BW, we observed a significant larger amplitude in alpha band in the MD group than that in the HC group (*t*_(60)_ = 2.58, *p* = 0.012, *Cohen’s d* = 0.64).

**Microscale**

For FW velocity, we observed a significant group × time interaction in the alpha band (*F*_(1,60)_ = 4.21, *p* = 0.045, *η²p* = 0.07). Post hoc analyses revealed that in HC, alpha wave velocity significantly decreased during the second view session (*t*_(60)_ = –2.24, *p* = 0.029, Cohen’s *d* = –0.21). The interaction is shown in the **Figure 4B**.

**Macroscale**

For BW, we found significant *group* × *part* (*F*_(2,120)_ = 3.37, *p* = 0.038, *η²p* = 0.05) and *group* × *part* × *time* (*F*_(2,120)_ = 3.94, *p* = 0.022, *η²p* = 0.06) interactions in the theta band. Post hoc comparisons indicated that in MD, BW theta velocity continuously decreased over the course of the experiment’s session (Late-Middle: *t*_(60)_ = –3.50, *p_FDR_* = 0.003, Cohen’s *d* = –0.65). The interaction is shown in the **Figure 4E**.

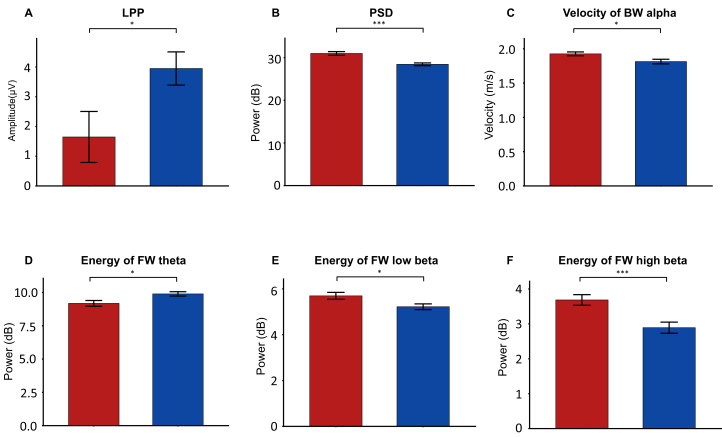

### Figure S1. Baseline neural abnormalities in response to single-shot negative stimuli. This figure illustrates the “baseline neural state” differences between Major Depressive (MD) Disorder patients (red bars) and Healthy Controls (HCs) (blue bars) during the first viewing of negative images, prior to any repetition effects. A. *Event-Related Potentials*. MD patients exhibit significantly reduced frontal Late Positive Potential (LPP) amplitudes compared to HCs, consistent with emotional blunting during initial processing. B. *Oscillatory Power*. MD patients show globally elevated Power Spectral Density (PSD) in the theta band, reflecting aberrant baseline cortical arousal. C. *Traveling Wave Velocity*. Disrupted top-down dynamics are evident in MD patients, characterized by paradoxically accelerated Backward (BW) Alpha velocity. D–F. *Traveling Wave Energy*. Large-scale forward propagation is systematically altered in MD, manifesting as (D) weakened Forward (FW) Theta energy, alongside (E) elevated FW Low-Beta and (F) FW High-Beta energy. These static signatures indicate that MD patients enter the task with an intrinsically disordered network state marked by high-frequency hyper-connectivity and low-frequency deficits. Error bars represent standard error. Significance markers: *** *p* < 0.001, * *p* < 0.05.

### Table S1. The mean and standard deviation of behavioral data

| Group | Part | *M*±*SD* |
| --- | --- | --- |
| MD | Early | 3.81±1.28 |
| MD | Middle | 3.73±1.35 |
| MD | Late | 3.73±1.46 |
| HC | Early | 3.03±1.14 |
| HC | Middle | 3.13±1.31 |
| HC | Late | 2.96±1.27 |

MD: major depression, HC: healthy controls, M: mean, SD: standard deviation

### Table S2. The mean and standard deviation of LPP.

| Group | Time | Part | *M*±*SD* |
| --- | --- | --- | --- |
| MD | First | Early | -0.33±4.97 |
| MD | First | Middle | 2.12±5.63 |
| MD | First | Late | 3.16±5.30 |
| MD | Second | Early | 1.64±4.80 |
| MD | Second | Middle | 1.59±5.00 |
| MD | Second | Late | 2.20±4.13 |
| HC | First | Early | 3.38±3.84 |
| HC | First | Middle | 4.20±3.41 |
| HC | First | Late | 4.27±3.84 |
| HC | Second | Early | 1.18±2.88 |
| HC | Second | Middle | 1.40±2.83 |
| HC | Second | Late | 0.02±1.87 |

LPP: late positive potential

### Table S3. The mean and standard deviation of N2.

| Group | Time | Part | *M*±*SD* |
| --- | --- | --- | --- |
| MD | First | Early | -7.85±5.88 |
| MD | First | Middle | -6.95±5.58 |
| MD | First | Late | -6.26±6.39 |
| MD | Second | Early | -5.18±5.28 |
| MD | Second | Middle | -3.79±6.67 |
| MD | Second | Late | -4.59±5.41 |
| HC | First | Early | -8.80±5.73 |
| HC | First | Middle | -7.98±5.87 |
| HC | First | Late | -8.35±5.99 |
| HC | Second | Early | -7.21±5.16 |
| HC | Second | Middle | -6.13±5.37 |
| HC | Second | Late | -6.84±5.86 |

### Table S4. The mean and standard deviation of PSD.

| Group | Time | Part | *M*±*SD* |
| --- | --- | --- | --- |
| MD | First | Early | 31.07±2.35 |
| MD | First | Middle | 30.81±2.20 |
| MD | First | Late | 31.08±2.40 |
| MD | Second | Early | 31.41±2.40 |
| MD | Second | Middle | 31.12±2.35 |
| MD | Second | Late | 31.06±2.52 |
| HC | First | Early | 28.34±1.81 |
| HC | First | Middle | 28.43±1.92 |
| HC | First | Late | 28.54±1.96 |
| HC | Second | Early | 28.03±2.02 |
| HC | Second | Middle | 28.07±1.94 |
| HC | Second | Late | 28.25±2.07 |

### Table S5. The mean and standard deviation of FW energy on theta band.

| Group | Time | Part | *M*±*SD* |
| --- | --- | --- | --- |
| MD | First | Early | 9.24±1.25 |
| MD | First | Middle | 9.16±1.51 |
| MD | First | Late | 9.15±1.44 |
| MD | Second | Early | 9.16±1.17 |
| MD | Second | Middle | 9.03±1.24 |
| MD | Second | Late | 8.98±1.23 |
| HC | First | Early | 9.73±1.01 |
| HC | First | Middle | 10.00±1.10 |
| HC | First | Late | 9.95±1.22 |
| HC | Second | Early | 9.80±1.12 |
| HC | Second | Middle | 9.30±1.15 |
| HC | Second | Late | 9.45±1.22 |

### Table S6. The mean and standard deviation of FW energy on alpha band.

| Group | Time | Part | *M*±*SD* |
| --- | --- | --- | --- |
| MD | First | Early | 7.52±0.85 |
| MD | First | Middle | 7.73±0.91 |
| MD | First | Late | 7.68±1.11 |
| MD | Second | Early | 7.63±1.01 |
| MD | Second | Middle | 7.52±0.71 |
| MD | Second | Late | 7.85±0.89 |
| HC | First | Early | 7.82±0.69 |
| HC | First | Middle | 7.85±0.95 |
| HC | First | Late | 7.99±0.77 |
| HC | Second | Early | 7.91±0.94 |
| HC | Second | Middle | 8.17±0.90 |
| HC | Second | Late | 8.02±0.83 |

### Table S7. The mean and standard deviation of FW energy on low beta band.

| Group | Time | Part | *M*±*SD* |
| --- | --- | --- | --- |
| MD | First | Early | 5.70±0.84 |
| MD | First | Middle | 5.69±1.01 |
| MD | First | Late | 5.69±1.03 |
| MD | Second | Early | 5.90±0.94 |
| MD | Second | Middle | 5.76±0.84 |
| MD | Second | Late | 5.74±0.88 |
| HC | First | Early | 5.37±0.83 |
| HC | First | Middle | 5.18±0.90 |
| HC | First | Late | 5.11±0.89 |
| HC | Second | Early | 5.37±1.01 |
| HC | Second | Middle | 5.60±0.75 |
| HC | Second | Late | 5.61±0.74 |

### Table S8. The mean and standard deviation of FW energy on high beta band.

| Group | Time | Part | *M*±*SD* |
| --- | --- | --- | --- |
| MD | First | Early | 3.87±0.90 |
| MD | First | Middle | 3.66±1.02 |
| MD | First | Late | 3.53±1.17 |
| MD | Second | Early | 3.93±1.29 |
| MD | Second | Middle | 3.77±1.21 |
| MD | Second | Late | 3.71±1.49 |
| HC | First | Early | 2.99±0.97 |
| HC | First | Middle | 2.86±0.98 |
| HC | First | Late | 2.83±1.23 |
| HC | Second | Early | 3.24±1.05 |
| HC | Second | Middle | 3.07±1.07 |
| HC | Second | Late | 3.00±1.17 |

### Table S9. The mean and standard deviation of BW energy on theta band.

| Group | Time | Part | *M*±*SD* |
| --- | --- | --- | --- |
| MD | First | Early | 5.63±2.40 |
| MD | First | Middle | 5.62±1.97 |
| MD | First | Late | 5.36±2.23 |
| MD | Second | Early | 5.62±2.18 |
| MD | Second | Middle | 5.63±2.49 |
| MD | Second | Late | 5.38±2.47 |
| HC | First | Early | 6.03±1.44 |
| HC | First | Middle | 6.03±1.41 |
| HC | First | Late | 6.00±1.34 |
| HC | Second | Early | 5.79±1.41 |
| HC | Second | Middle | 5.72±1.17 |
| HC | Second | Late | 5.89±1.47 |

### Table S10. The mean and standard deviation of BW energy on alpha band.

| Group | Time | Part | *M*±*SD* |
| --- | --- | --- | --- |
| MD | First | Early | 3.79±2.25 |
| MD | First | Middle | 4.07±1.93 |
| MD | First | Late | 3.72±2.22 |
| MD | Second | Early | 3.54±2.27 |
| MD | Second | Middle | 3.89±2.56 |
| MD | Second | Late | 3.44±2.28 |
| HC | First | Early | 4.29±0.94 |
| HC | First | Middle | 4.22±1.06 |
| HC | First | Late | 4.52±1.06 |
| HC | Second | Early | 4.01±1.01 |
| HC | Second | Middle | 4.40±1.04 |
| HC | Second | Late | 4.50±1.07 |

### Table S11. The mean and standard deviation of BW energy on low beta band.

| Group | Time | Part | *M*±*SD* |
| --- | --- | --- | --- |
| MD | First | Early | 1.47±2.06 |
| MD | First | Middle | 1.75±1.80 |
| MD | First | Late | 1.91±2.08 |
| MD | Second | Early | 1.69±2.02 |
| MD | Second | Middle | 1.96±2.25 |
| MD | Second | Late | 1.52±2.15 |
| HC | First | Early | 1.53±0.97 |
| HC | First | Middle | 1.61±1.02 |
| HC | First | Late | 1.20±1.03 |
| HC | Second | Early | 1.60±0.98 |
| HC | Second | Middle | 1.68±1.03 |
| HC | Second | Late | 1.58±1.18 |

### Table S12. The mean and standard deviation of BW energy on high beta band.

| Group | Time | Part | *M*±*SD* |
| --- | --- | --- | --- |
| MD | First | Early | -0.69±2.37 |
| MD | First | Middle | -0.58±2.00 |
| MD | First | Late | -0.72±2.03 |
| MD | Second | Early | -0.78±2.13 |
| MD | Second | Middle | -0.61±2.40 |
| MD | Second | Late | -0.69±2.17 |
| HC | First | Early | -1.04±1.07 |
| HC | First | Middle | -1.15±1.05 |
| HC | First | Late | -1.35±0.99 |
| HC | Second | Early | -1.01±0.99 |
| HC | Second | Middle | -1.04±1.00 |
| HC | Second | Late | -1.08±1.03 |

### Table S13. The mean and standard deviation of FW velocity on theta band.

| Group | Time | Part | M±SD |
| --- | --- | --- | --- |
| MD | First | Early | 1.05±0.10 |
| MD | First | Middle | 1.04±0.10 |
| MD | First | Late | 1.02±0.12 |
| MD | Second | Early | 1.03±0.10 |
| MD | Second | Middle | 1.04±0.10 |
| MD | Second | Late | 1.03±0.10 |
| HC | First | Early | 1.03±0.11 |
| HC | First | Middle | 1.05±0.10 |
| HC | First | Late | 1.04±0.08 |
| HC | Second | Early | 1.03±0.09 |
| HC | Second | Middle | 1.03±0.09 |
| HC | Second | Late | 1.01±0.09 |

### Table S14. The mean and standard deviation of FW velocity on alpha band.

| Group | Time | Part | M±SD |
| --- | --- | --- | --- |
| MD | First | Early | 2.01±0.15 |
| MD | First | Middle | 1.95±0.22 |
| MD | First | Late | 1.95±0.19 |
| MD | Second | Early | 1.99±0.18 |
| MD | Second | Middle | 1.96±0.16 |
| MD | Second | Late | 2.00±0.13 |
| HC | First | Early | 2.03±0.13 |
| HC | First | Middle | 1.98±0.17 |
| HC | First | Late | 2.00±0.16 |
| HC | Second | Early | 1.95±0.17 |
| HC | Second | Middle | 1.97±0.14 |
| HC | Second | Late | 1.97±0.16 |

### Table S15. The mean and standard deviation of FW velocity on low beta band.

| Group | Time | Part | M±SD |
| --- | --- | --- | --- |
| MD | First | Early | 3.16±0.26 |
| MD | First | Middle | 3.13±0.23 |
| MD | First | Late | 3.16±0.29 |
| MD | Second | Early | 3.28±0.19 |
| MD | Second | Middle | 3.17±0.27 |
| MD | Second | Late | 3.14±0.30 |
| HC | First | Early | 3.22±0.24 |
| HC | First | Middle | 3.23±0.22 |
| HC | First | Late | 3.23±0.19 |
| HC | Second | Early | 3.19±0.28 |
| HC | Second | Middle | 3.19±0.19 |
| HC | Second | Late | 3.25±0.20 |

### Table S16. The mean and standard deviation of FW velocity on high beta band.

| Group | Time | Part | M±SD |
| --- | --- | --- | --- |
| MD | First | Early | 4.67±0.44 |
| MD | First | Middle | 4.76±0.40 |
| MD | First | Late | 4.65±0.40 |
| MD | Second | Early | 4.69±0.35 |
| MD | Second | Middle | 4.77±0.35 |
| MD | Second | Late | 4.66±0.31 |
| HC | First | Early | 4.70±0.27 |
| HC | First | Middle | 4.75±0.30 |
| HC | First | Late | 4.78±0.29 |
| HC | Second | Early | 4.81±0.28 |
| HC | Second | Middle | 4.74±0.26 |
| HC | Second | Late | 4.77±0.28 |

### Table S17. The mean and standard deviation of BW velocity on theta band.

| Group | Time | Part | M±SD |
| --- | --- | --- | --- |
| MD | First | Early | 0.99±0.11 |
| MD | First | Middle | 0.99±0.12 |
| MD | First | Late | 0.99±0.13 |
| MD | Second | Early | 1.03±0.10 |
| MD | Second | Middle | 0.99±0.09 |
| MD | Second | Late | 0.93±0.17 |
| HC | First | Early | 0.98±0.12 |
| HC | First | Middle | 0.96±0.14 |
| HC | First | Late | 0.99±0.14 |
| HC | Second | Early | 0.95±0.13 |
| HC | Second | Middle | 1.00±0.11 |
| HC | Second | Late | 1.00±0.11 |

### Table S18. The mean and standard deviation of BW velocity on alpha band.

| Group | Time | Part | M±SD |
| --- | --- | --- | --- |
| MD | First | Early | 1.91±0.18 |
| MD | First | Middle | 1.94±0.19 |
| MD | First | Late | 1.94±0.17 |
| MD | Second | Early | 1.90±0.21 |
| MD | Second | Middle | 1.93±0.23 |
| MD | Second | Late | 1.88±0.26 |
| HC | First | Early | 1.80±0.22 |
| HC | First | Middle | 1.78±0.24 |
| HC | First | Late | 1.86±0.23 |
| HC | Second | Early | 1.79±0.23 |
| HC | Second | Middle | 1.83±0.22 |
| HC | Second | Late | 1.87±0.15 |

### Table S19. The mean and standard deviation of BW velocity on low beta band.

| Group | Time | Part | M±SD |
| --- | --- | --- | --- |
| MD | First | Early | 3.03±0.31 |
| MD | First | Middle | 3.00±0.38 |
| MD | First | Late | 3.01±0.34 |
| MD | Second | Early | 3.05±0.34 |
| MD | Second | Middle | 3.01±0.28 |
| MD | Second | Late | 2.96±0.39 |
| HC | First | Early | 3.09±0.24 |
| HC | First | Middle | 3.10±0.27 |
| HC | First | Late | 3.07±0.25 |
| HC | Second | Early | 3.06±0.28 |
| HC | Second | Middle | 3.10±0.29 |
| HC | Second | Late | 3.12±0.25 |

### Table S20. The mean and standard deviation of BW velocity on high beta band.

| Group | Time | Part | M±SD |
| --- | --- | --- | --- |
| MD | First | Early | 4.46±0.38 |
| MD | First | Middle | 4.53±0.42 |
| MD | First | Late | 4.34±0.59 |
| MD | Second | Early | 4.45±0.52 |
| MD | Second | Middle | 4.46±0.50 |
| MD | Second | Late | 4.49±0.49 |
| HC | First | Early | 4.46±0.42 |
| HC | First | Middle | 4.67±0.32 |
| HC | First | Late | 4.49±0.32 |
| HC | Second | Early | 4.51±0.41 |
| HC | Second | Middle | 4.53±0.37 |
| HC | Second | Late | 4.52±0.30 |
